# Deficient Cardiolipin Remodeling Alters Muscle Fiber Composition and Neuromuscular Connectivity in Barth Syndrome

**DOI:** 10.1101/2025.10.31.685856

**Authors:** Catalina Matias, Paige L. Snider, Elizabeth A. Sierra Potchanant, Joshua R. Huot, Rahul Raghav, Michael T. Chin, Simon J. Conway, Jeffrey J. Brault

**Affiliations:** Indiana Center for Musculoskeletal Health, Indiana University School of Medicine, Indianapolis, IN 46202, USA; Department of Anatomy, Cell Biology C Physiology, Indiana University School of Medicine, Indianapolis, IN 46202, USA; Herman B. Wells Center for Pediatric Research, Indiana University School of Medicine, Indianapolis, IN 46202, USA; Melvin and Bren Simon Comprehensive Cancer Center, Indiana University School of Medicine, Indianapolis, IN 46202, USA; Molecular Cardiology Research Institute, Tufts Medical Center, Boston, MA 02111, USA

## Abstract

**Background:** Barth syndrome (BTHS) is a rare X-linked mitochondrial disorder caused by mutations in the *TAFAZZIN* gene, which disrupts cardiolipin (CL) remodeling and mitochondrial function. While cardiac manifestations of BTHS are well characterized, the mechanisms underlying skeletal muscle weakness and fatigability are poorly understood.

**Methods:** We investigated neuromuscular and mitochondrial alterations in a novel murine model (Taz^PM^) carrying a patient-derived D75H point mutation in *Tafazzin*. This mutation preserves protein abundance but abolishes enzymatic activity. Skeletal muscle function was assessed via weightlifting and hanging tests. Muscle fiber composition and neuromuscular junction (NMJ) integrity were evaluated using immunofluorescence, western blotting, and in vivo electrophysiology. Mitochondrial morphology was examined by transmission electron microscopy, and bioenergetics were quantified using ultra-performance liquid chromatography. Stress signaling was assessed by western blotting.

**Results:** Male Taz^PM^ mice exhibited elevated monolysocardiolipin and reduced mature CL levels, confirming deficient transacylase activity. These mice exhibited lower muscle strength and endurance, smaller muscle fibers of all types, and a shift toward fast-twitch type 2B fibers, which are more susceptible to fatigue. Electrophysiological analysis revealed a 60% reduction in motor unit number and an increase in average single motor unit potential, indicating motor neuron remodeling. NMJ protein analysis showed decreased MUSK and DOK7 and increased CHRNA1, suggesting impaired NMJ integrity. Despite mitochondrial structural abnormalities and reduced expression of key mitochondrial proteins (NDUFB8, MCU, TMEM65), resting ATP, phosphocreatine, and adenine nucleotide ratios were unchanged in both glycolytic and oxidative muscles. However, stress signaling pathways were markedly activated, including phosphorylation of eIF2α, increased CHOP, DELE1, p53 expression, and altered Wnt/β-catenin signaling components.

**Conclusions:** Deficiency of Tafazzin enzymatic activity in skeletal muscle is sufficient to result in widespread neuromuscular remodeling, including fiber size/type shifts, motor unit loss, NMJ dysregulation, and stress pathway activation, without overt energetic failure at rest. These findings suggest that myopathy in BTHS arises not solely from mitochondrial ATP insufficiency but rather from cumulative structural and signaling disruptions.

## Introduction

TAFAZZIN is a mitochondrial transacylase enzyme that can serve as a protein scaffold [1, 2] and is essential for mitochondrial inner membrane stabilization and phospholipid remodeling. Specifically, TAFAZZIN catalyzes the conversion of immature monolysocardiolipin (MLCL) to mature tetralinoleoyl cardiolipin (CL). Additionally, the TAFAZZIN protein itself can sense inner mitochondrial membrane curvature [3] and binds protein complexes containing ADP/ATP-carrier/ATP-synthase and prohibitin/Dnajc19 [2, 4], thereby ensuring standard mitochondrial ultrastructure, electron transport chain complex alignment, and effective oxidative phosphorylation (OxPhos). Thus, when either TAFAZZIN protein or CL remodeling is disrupted, mitochondrial function is impaired, often leading to reduced maximal oxidative ATP-producing capacity, oxidative stress, and defective mitophagy [5–7]. However, barring the heart, the specific effects of non-functional TAFAZZIN and reduced CL in mitochondrial-enriched tissues remain unclear.

Loss-of-function mutations in the *TAFAZZIN* gene cause Barth syndrome (BTHS), a rare X-linked mitochondrial disorder primarily affecting males. More than 200 distinct mutations have been identified [S1], resulting in enzymatically inactive proteins or complete protein loss. Clinically, BTHS is characterized by growth restriction, cardiomyopathy, neutropenia, and skeletal myopathy [8]. Skeletal muscle abnormalities are evident from birth, presenting as hypotonia and delayed motor development. As affected individuals age, they experience difficulty with fine motor skills, profound fatigue, and muscle weakness [8–12], which are reported as the most debilitating symptoms impacting their daily living and quality of life [13]. Importantly, these skeletal muscle symptoms persist even after cardiac transplantation [14], indicating that muscle defects are not solely a secondary effect of cardiac insufficiency.

Despite the clinical significance of skeletal myopathy in BTHS, the underlying neuromuscular mechanisms remain poorly understood. Previous *Tafazzin* knockdown or knockout mouse models have shown muscle weakness [6, 15], increased fatigability [15–16], and muscle atrophy [16]. However, direct assessments of key factors that impact muscle weakness, such as comprehensive cellular energetics [17], muscle fiber typing/sizing, and neuromuscular connectivity, have not been reported.

To address this gap, we used a novel murine model of BTHS with a D75H substitution in *TAFAZZIN*, which mirrors the mutation of a young patient with classic BTHS symptoms [18]. This mutation is within the transacylase motif and results in normal protein levels in various organs and skeletal muscle [18]. However, enzyme activity is severely impaired by this mutation, as evidenced by a markedly increased MLCL/CL ratio in the blood [18], an established diagnostic marker for BTHS [19]. These mice exhibit progressive cardiomyopathy [18], decreased cardiac ATP levels as adults [18], testicular infertility [S3], and neutropenia [20].

The purpose of this study is to investigate mechanisms underlying skeletal muscle weakness and fatigability in a BTHS patient-derived D75H point mutant knock-in (*Taz^PM^*) mouse. We focus on major factors known to regulate muscle force production, including muscle fiber size and fiber type, cellular energetics, motor unit connectivity, along with mitochondrial homeostasis and stress responses. Understanding how these mechanisms are disrupted by deficiency of TAFAZZIN function may enable the development of new therapies targeting skeletal muscle, which remains a major unmet need in BTHS.

## Methods

### Mice

Wild-type (*wt*) and littermate *Tafazzin* (MGI:109626) D75H point mutant knockin (*Taz^PM^*) mice were generated by breeding heterozygous *Taz^wt^*/*Taz^PM^* females with *wt* males, as described previously [18]. Because BTHS is X-linked, predominantly affects males, and our *Taz^PM^* female mice are not overtly symptomatic, only male mice were examined in this study. All experimental mice were 4-7 months old, and *Taz^PM^* mice were always age-matched with their *wt* littermates. Genotyping was performed by PCR as described previously [18]. All animal procedures were approved by the Indiana University School of Medicine Animal Care and Use Committee (IACUC #23095).

### MLCL and CL Quantification

Adult mitochondria were isolated from gastrocnemius muscles following trypsin digestion and sucrose extraction, and then spotted onto Whatman protein saver cards, as described [18]. Dried extracts were eluted with a mixture of acetonitrile/isopropanol/water. Lipid extracts were separated by reverse-phase HPLC in ZORBAX Eclipse Plus C18 2.1 × 50 mm, 1.8 μm particles (Agilent, Santa Clara) with solvent A (Acetonitrile/Water [60:40], 10 mM ammonium formate, 0.1% formic acid) and solvent B (Isopropanol/Acetonitrile [90:10], 10 mM ammonium formate, 0.1% formic acid) at 55 °C and analyzed via 6495A Triple Quadrupole LC/MS System (Agilent) [18].

### Muscle Contractile Function Testing

Mice underwent a weightlifting test using steel wool pads attached to incrementally heavier metal rings. Mice were suspended by their tails above the pads, which they instinctively grasped. Once a secure grip was established, mice were lifted along with the weighted pad and tested for the ability to hold the weight for 3 seconds. After each trial, mice rested for 1 minute before the next test, with weight increased by 8 g per trial (range: 16.9-58.1 g). The maximum weight held for 3 seconds was recorded.

Neuromuscular function was further assessed using a hanging bar test. Mice were allowed to grasp an elevated horizontal bar with their forepaws. The duration mice could support their body weight before falling was recorded. The body weight of each mouse was measured before the exercise.

### Immunoffuorescence Staining

Muscle fiber typing was performed as we have done previously [21]. Procedure details for muscle processing and immunostaining are provided in Supplemental Method S1.

Antibody information is provided in Table S1. Images were analyzed for fiber type-specific cross-sectional area using the freely available software QuantiMus [S4].

### In vivo Electrophysiology

Mice were anesthetized with inhaled 2% isoflurane in oxygen, and sciatic nerve of the left limb was stimulated with two 28-gauge electrodes (Natus Neurology); a duo shielded ring electrode was used for recording, and a ground electrode was placed on the animal’s tail as done previously [S5]. Baseline-to-peak and peak-to-peak compound muscle action potential (CMAP) responses were recorded utilizing supramaximal stimulations (constant current intensity: <10 mA; pulse duration: 0.1 ms). Single motor unit potential (SMUP) size was determined using an incremental stimulation technique. Incremental responses were obtained by submaximal stimulation of the sciatic nerve until a stable, minimal all-or-none response occurred. Ten successive SMUP increments were recorded and averaged. Motor unit number estimation (MUNE) was calculated by the following: CMAP amplitude (peak-to-peak)/average SMUP (peak-to-peak).

### Western Blot

Gastrocnemius muscles were homogenized in RIPA protein lysis buffer (MilliporeSigma) and ∼20–30 μg sample/lane resolved using SDS-PAGE (Bio-Rad) and transferred to PVDF Western Blotting Membranes, as described [18]. Following blocking in Blotting-Grade Blocker (Bio-Rad), blots were probed with primary (Table S1) and secondary antibodies (Table S2). Immunoreactive protein band signals were detected via ECL Western Blotting Detection (Amersham, UK) with species-appropriate peroxidase-conjugated secondary antibodies (Bio-Rad and Jackson ImmunoResearch, West Grove, PA, USA). To verify equal loading, all blots were subsequently stripped (Fisher Stripping Buffer for 15 min at 37°C), washed, re-blocked, and then probed with housekeeping GAPDH (Table S1) to monitor protein integrity and loading. X-ray films of varying exposures were scanned for each antibody, and the densitometric signal intensities were quantified.

### Transmission Electron Microscopy

Gastrocnemius muscles were isolated and fixed in 2% paraformaldehyde and 2% glutaraldehyde fixative in 0.1 M cacodylate buffer (pH 7.2) overnight, processed, and imaged on a Phillips EM 400 microscope (via IU EM Core) as described [18]. Mitochondrial morphology was assessed in a manner blinded to genotype and analyzed using ImageJ (version 1.54j).

### Ultra Performance Liquid Chromatography

Tissue collection, extraction of analytes, and analysis of nucleotides, phosphocreatine, and NAD+/NADH were performed as described previously [18, S6]. Details are given in Supplemental Information.

### Statistics

Data presented as mean ± standard deviation, with p<0.05 considered significant. Statistical analysis was performed via Prism software version 5.02 (GraphPad Software). Comparisons between the two groups, *wt* and *Taz^PM^*, were performed by unpaired t-test in all cases except for weightlifting, where the Mann-Whitney test was used because the data were ordinal rather than continuous. Comparisons of lipid species, fiber type distribution, and fiber type-specific area were performed by 2-way ANOVA followed by Sidak post-hoc test.

## Results

To confirm the loss of muscle TAFAZZIN enzymatic acyl chain remodeling activity in *Taz^PM^*, we measured immature MLCL and mature CL species levels in mitochondria isolated from gastrocnemius muscles. In *Taz^PM^*, MLCL(18:2)3 was decreased, while all other MLCL species measured, MLCL(18:1)2(16:1), (18:1)(16:0)2, (18:1)2(16:0), and (18:1)3, were significantly increased (**Figure 1a**). Consequently, total MLCL was 7-fold greater in *Taz^PM^* versus *wt*. In *Taz^PM^*, CL(18:2)4, CL(18:2)3(16:1), and (18:1)2(18:2)(16:1) species were dramatically decreased, while CL(18:1)2(16:0)2, (18:1)3(16:0), and (18:1)4 species were increased (**Figure 1b**). Since CL(18:2)4 is overwhelmingly the predominant CL mature species in *wt*, total CL was 5-fold less in *Taz^PM^*. Combined, this resulted in a 36-fold greater MLCL:CL ratio in *Taz^PM^* muscle mitochondria (**Figure 1c**), the gold-standard diagnostic biomarker of BTHS [19]. As described in our previous study [18], adult *Taz^PM^* mice were smaller than their *wt* littermates (**Figure 1d**). To test for neuromuscular function, mice were evaluated for their maximal lifting ability and duration of bar hanging. As expected, *Taz^PM^* mice exhibited decreased maximal weightlifting capacity (**Figure 1e**) and hung from a bar for a shorter duration (**Figure 1f**). Therefore, the *Taz^PM^* mice faithfully replicate muscle weakness and growth restriction observed in BTHS patients.

**Figure 1.**
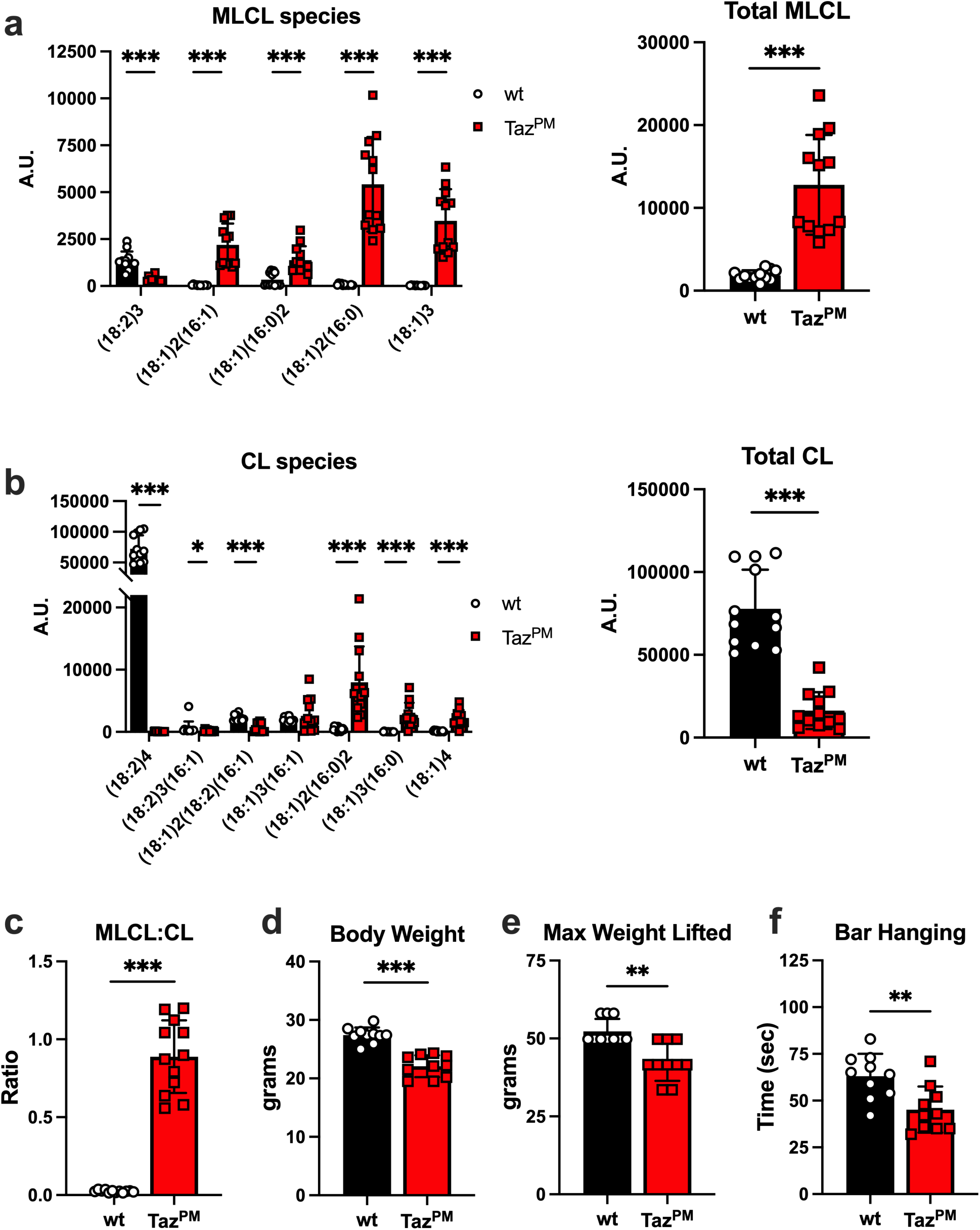
Cardiolipin species and skeletal muscle phenotype of *Taz^PM^*. (a) Mass spectrometry quantification of individual monolysocardiolipin (MLCL) species and total MLCL (sum of individual species) in isolated mitochondria from age-matched *wt* and *Taz^PM^* gastrocnemius muscles. n=12 per genotype. (b) Mass spectrometry quantification of individual cardiolipin (CL) species and total CL (sum of individual species) (c) Calculation of total MLCL to total CL ratio. (d) Body weight n=10 per genotype, (e) maximal weight lifted per mouse, and (f) duration of hanging from a high bar. n=10 per genotype. **p<0.05, **p<0.01, ***p<0.001

Skeletal muscle strength is associated with myosin heavy chain expression, predominantly due to differences in the cross-sectional area (CSA) of different fiber types.

Therefore, we next tested whether *Taz^PM^* induces a shift in the proportion and/or size of muscle fibers expressing different myosin heavy chains. Using immunofluorescence for myosin heavy chain and laminin, we were able to discern the adult myosin heavy chains type 2A, type 2X, and type 2B but not type 1 in tibialis anterior muscles (**Figure 2a**).

**Figure 2.**
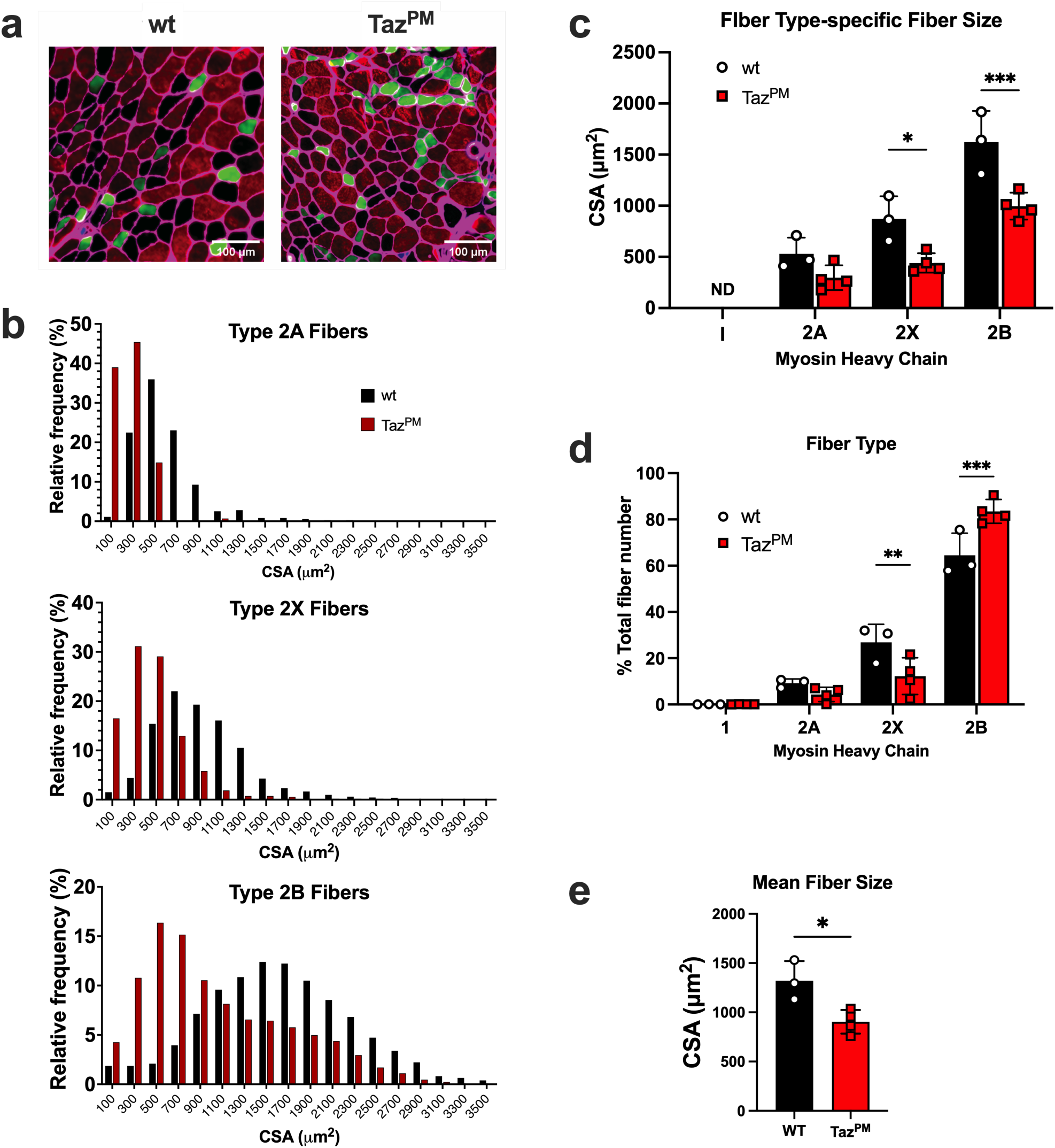
Skeletal muscle fibers are smaller and exhibit a shift toward a faster myosin heavy chain in the *Taz^PM^* mice. (a) Immunofluorescence for laminin (magenta), type 1 myosin heavy chain (MHC)(blue), type 2A MHC (green), and type 2B MHC (red) was performed on age-matched *wt* and *Taz^PM^* tibialis anterior muscle cross-sections. Unstained muscle fibers are type 2x MHC (black). (b) Frequency distribution of fiber size separated by MHC expression. (c) Mean fiber type-specific cross-sectional area (CSA) fiber size. ND = none detected. (d) The proportion of fibers staining positive for each MHC. (e) Mean area of all fibers. n=3-4 per genotype. *p<0.05, **p<0.01, ***p<0.001.

Interestingly, whereas the fiber type staining of *wt* appeared in a typical checkboard pattern, in *Taz^PM^* the type 2A fibers appeared abnormally grouped. The frequency distribution for fiber size was leftward shifted for all fiber types detected, suggesting all fibers become smaller, not just a subset (**Figure 2b**). Accordingly, the mean fiber area for type 2X and type 2B fibers was significantly smaller in *Taz^PM^* versus *wt* (**Figure 2c**). There was also a significant change in fiber type with a greater proportion of larger type 2B fibers and a lower proportion of 2X fibers in *Taz^PM^* (**Figure 2d**). The mean muscle fiber size of all fibers was significantly smaller in *Taz^PM^* (**Figure 2e**). Interestingly, we also observed an increased proportion of centrally nucleated fibers (**Figure S1**), suggesting active or recent fiber remodeling.

Muscle fiber type grouping, as in *Taz^PM^*, is a classic indicator of muscle fiber denervation and reinnervation by a neighboring, surviving motor neuron [S7]. Therefore, motor unit number might be expected to decrease in *Taz^PM^*, as seen, for example, in amyotrophic lateral sclerosis [S8]. To test this possibility, we performed *in vivo* electrophysiology of the triceps surae (gastrocnemius, plantaris, and soleus) muscles. In *Taz^PM^*, SMUP was increased 2.8-fold (**Figure 3a**), and MUNE was decreased by 60% (**Figure 3b**), indicating that motor unit connectivity was decreased in these mutant muscles.

**Figure 3.**
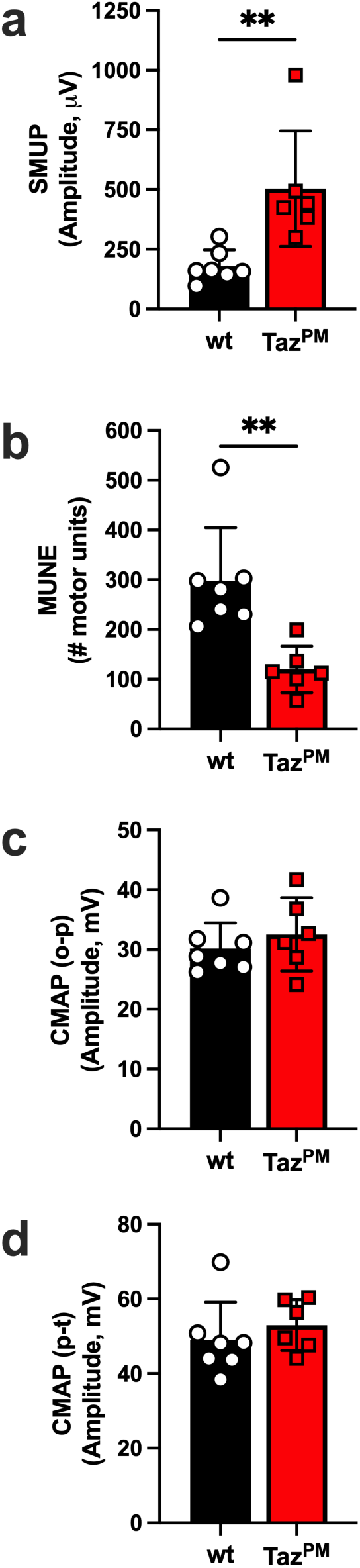
Motor unit number decreases in *Taz^PM^* hindlimb. (a) Single motor unit potential (SMUP), (b) motor unit number estimate (MUNE), (c) compound muscle action potential from onset to peak (CMAP (o-p)), and (d) compound muscle action potential from peak to trough (CMAP (p-t)) were measured in vivo from age-matched *wt* and *Taz^PM^* triceps surae muscles. n=6-7 per genotype. **p<0.01

However, the amplitudes of the CMAP in the onset-to-peak phase (CMAP(o-p)) and peak-to-trough phase (CMAP(p-t)) were not different between *wt* and *Taz^PM^* (**Figure 3c, d**), indicating there was no discernible difference in overall innervation of the hindlimb muscles, at least at maximal stimulating conditions. Taken together, this supports a loss of smaller motor units with larger motor units sustaining *Taz^PM^* innervation.

To better understand the potential mechanisms underlying changes in muscle innervation, we imaged the neuromuscular junction (NMJ) in *wt* (**Figure 4a**) and *Taz^PM^* (**Figure 4b**) gastrocnemius muscles via immunofluorescence. The vast majority of staining for Synapsin (found on the presynaptic terminus) and BTX (binds to the acetylocholine receptor (AChRs) in the postsynaptic terminus) was co-localized (**Figure 4a top G bottom, Figure 4b top**), as would be expected for proper functioning NMJs. However, there was evidence in *Taz^PM^* that Synapsin and BTX staining were not co-localized (**Figure 4b bottom**). To quantify possible differential protein expression on the whole muscle level, we performed western blotting for vital NMJ-related proteins (**Figure 4c**). On the presynaptic side of the NMJ, SNAP25 (essential for synaptic fusion [S9]), α tubulin, and β tubulin (both important for axonal transport and NMJ stability [S10]) are not different between *wt* and *Taz^PM^* (**Figure 4d**). In contrast, proteins of the postsynaptic side were differentially expressed. CHRNA1, a subunit of the AChR, was increased in *Taz^PM^* muscle (**Figure 4e**).

**Figure 4.**
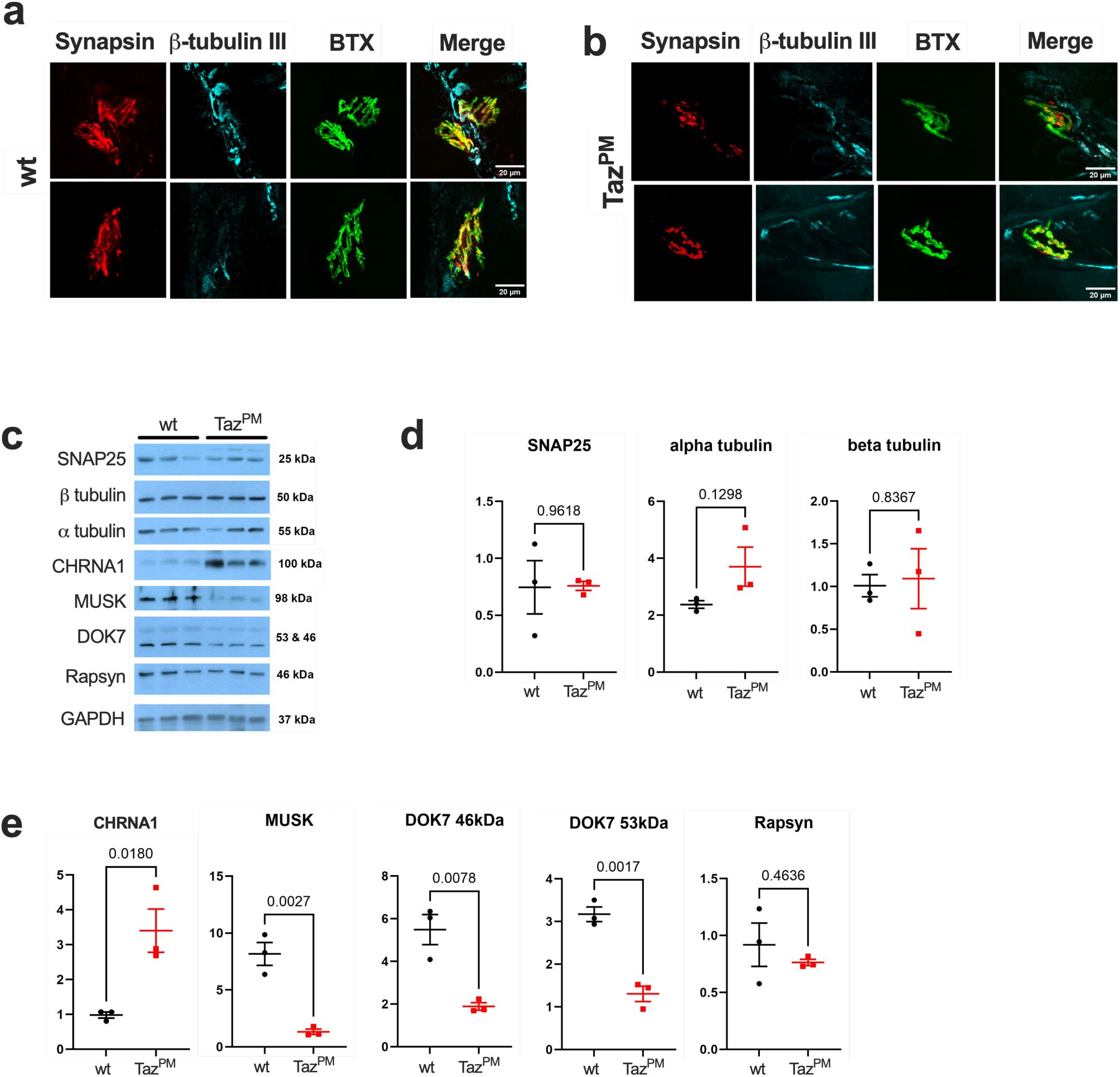
Proteins of the neuromuscular junction are dysregulated in muscle of *Taz^PM^*. Representative immunofluorescent imaging for Synapsin, β tubulin III, BTX, and merged images for (a) *wt* and (b) *Taz^PM^* gastrocnemius. (c) Western blot images for synaptosomal-associated protein-25 (SNAP25), α tubulin, β tubulin, cholinergic receptor nicotinic alpha 1 subunit (CHRNA1), muscle associated receptor tyrosine kinase (MUSK), dedicator of cytokinesis 7 (DOK7), Rapsyn, and glyceraldehyde 3-phosphate dehydrogenase (GAPDH). (d) Quantification of western blots for presynaptic proteins SNAP25, α tubulin, and β tubulin. (e) Quantification of the western blots for the postsynaptic proteins CHRNA1, MUSK, DOK7 46 kDa band, DOK7 53 kDa band, and Rapsyn. n=3 per genotype.

MUSK, a receptor tyrosine kinase [S11], and DOK7, which binds to and further activates MUSK [S12], were both decreased in *Taz^PM^* (**Figure 4e**). Rapsyn, which acts as a linker between the AChRs and the cytoskeleton [S13], was expressed to the same degree in both *wt* and *Taz^PM^* muscles (**Figure 4e**). Therefore, proteins associated with the NMJ are differentially regulated in *Taz^PM^* muscles, which supports aberrant remodeling of muscle innervation.

Loss of Tafazzin transacylase activity and consequent loss of CL result in mitochondrial structural defects and bioenergetics deficits in cardiac muscle [18] and cultured cells [22]. Therefore, we performed TEM to examine mitochondrial ultrastructure in the gastrocnemius muscle (**Figure 5a**). Mitochondria of *Taz^PM^* muscle were swollen, and cristae were decreased in number and appeared disorganized, suggestive of impaired ATP production, disorganized ETC complex arrangement, abnormal protein gradient maintenance, and/or ROS accumulation. To test whether mitochondrial protein expression was altered, we performed western blots for Citrate synthase (CS), Voltage dependent anion-selective channel 1 (VDAC1), Succinate dehydrogenase subunit A (SDHA, an enzyme found both in the TCA cycle and OXPHOS), ATP synthase F1 subunit alpha (a subunit of mitochondrial complex V), NDUFB8 (a subunit of mitochondrial complex I), Mitochondrial calcium uniporter (MCU), and Transmembrane protein 65 (TMEM65) (**Figure 5b**). CS, a commonly used marker of mitochondrial content in muscle, was statistically unchanged in *Taz^PM^* (**Figure 5c**). Similarly, mitochondrial VDAC1, SDHA, and ATP5a1 protein levels were not different between *wt* and *Taz^PM^*, indicating that mutant mitochondrial content was unaffected. However, NDUFB8, MCU, and TMEM65 were drastically reduced in *Taz^PM^* (**Figure 5d**). Therefore, *Taz^PM^* muscle exhibits selective, significant decreases in key regulators of mitochondrial Ca^2+^ flux (MCU, TMEM65) and at least one mitochondrial complex 1 protein (NDUFB8) but not in the mitochondrial complex V protein (ATP5a1).

**Figure 5.**
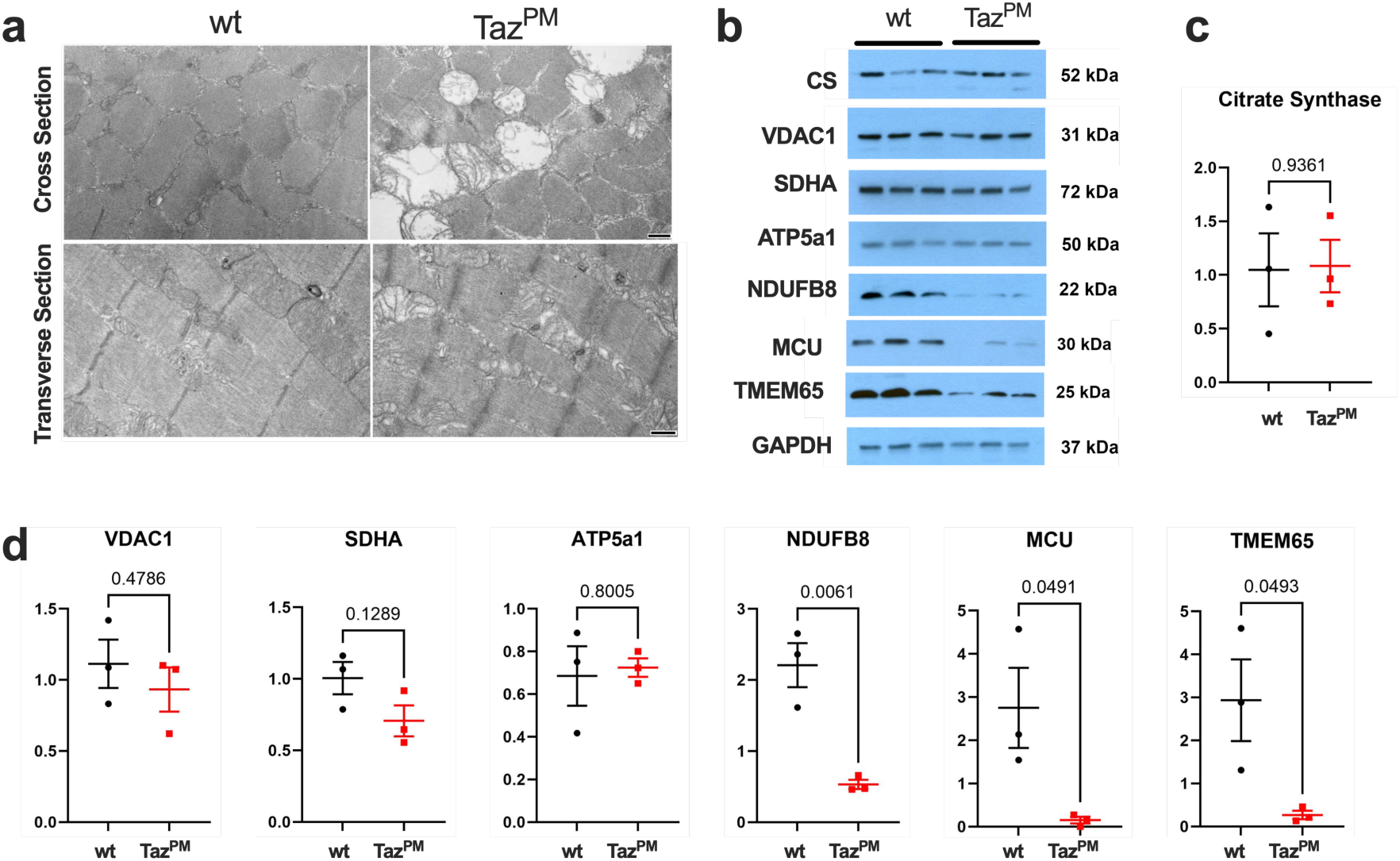
Mitochondrial morphology is deranged without consistent changes in mitochondrial protein content in muscle of *Taz^PM^* mice. (a) Representative TEM images of cross-section (Top) and longitudinal sections (Bottom) of *wt* (Left) and *Taz^PM^* (Right) gastrocnemius muscle. Scale bars Top = 800 nm, Bottom = 1 μm. (b) Western blots for mitochondrial proteins citrate synthase (CS), voltage dependent anion-selective channel 1 (VDAC1), succinate dehydrogenase (SDHA), ATP synthase F1 subunit alpha (ATP5a1), and NADH Ubiquinone oxidoreductase subunit B8 (NDUFB8), mitochondrial calcium uniporter (MCU), transmembrane protein 65 (TMEM65), and glyceraldehyde-3-phosphate dehydrogenase (GAPDH). (c) Quantification of citrate synthase blot. (d) Mitochondrial protein western blot quantification. n=3 per genotype.

Given that a major function of mitochondria is maintenance of the cellular energetic state and that muscle fibers and NMJ are highly energetic sites, we next examined key high-energy phosphate molecules. ATP, ADP, and AMP were all unchanged in gastrocnemius muscle of *Taz^PM^* (**Figure 6a**). The adenylate energy charge [S14], the ADP/ATP ratio (an indicator of free energy of ATP hydrolysis [23]), and the AMP/ATP ratio (known for activating AMP activated protein kinase [S14]) were also all unchanged in *Taz^PM^* (**Figure 6b**). The sum of adenine nucleotides (which is regulated by the balance between purine *de novo* synthesis and degradation [S15]), uric acid, an indicator of purine nucleotide (ATP or GTP) degradation [S16], (**Figure 6c**), and phosphocreatine (PCr), a high-energy phosphate reservoir that buffers ATP [23], (**Figure 6d**) were similarly unchanged (**Figure 6d**). Likewise, NAD+, NADH, and the NAD+/NADH ratio, indicators of the redox state and regulators of metabolism [S17], were not different between *Taz^PM^* and *wt* (**Figure 6e**). Therefore, despite our findings of major morphological changes in mutant muscle mitochondria, as well as decreased complex I protein NDUFB8 and mitochondrial Ca2+ regulating MCU and TMEM65 proteins, bioenergetics appear unchanged in the predominantly fast-twitch gastrocnemius muscle of *Taz^PM^*.

**Figure 6.**
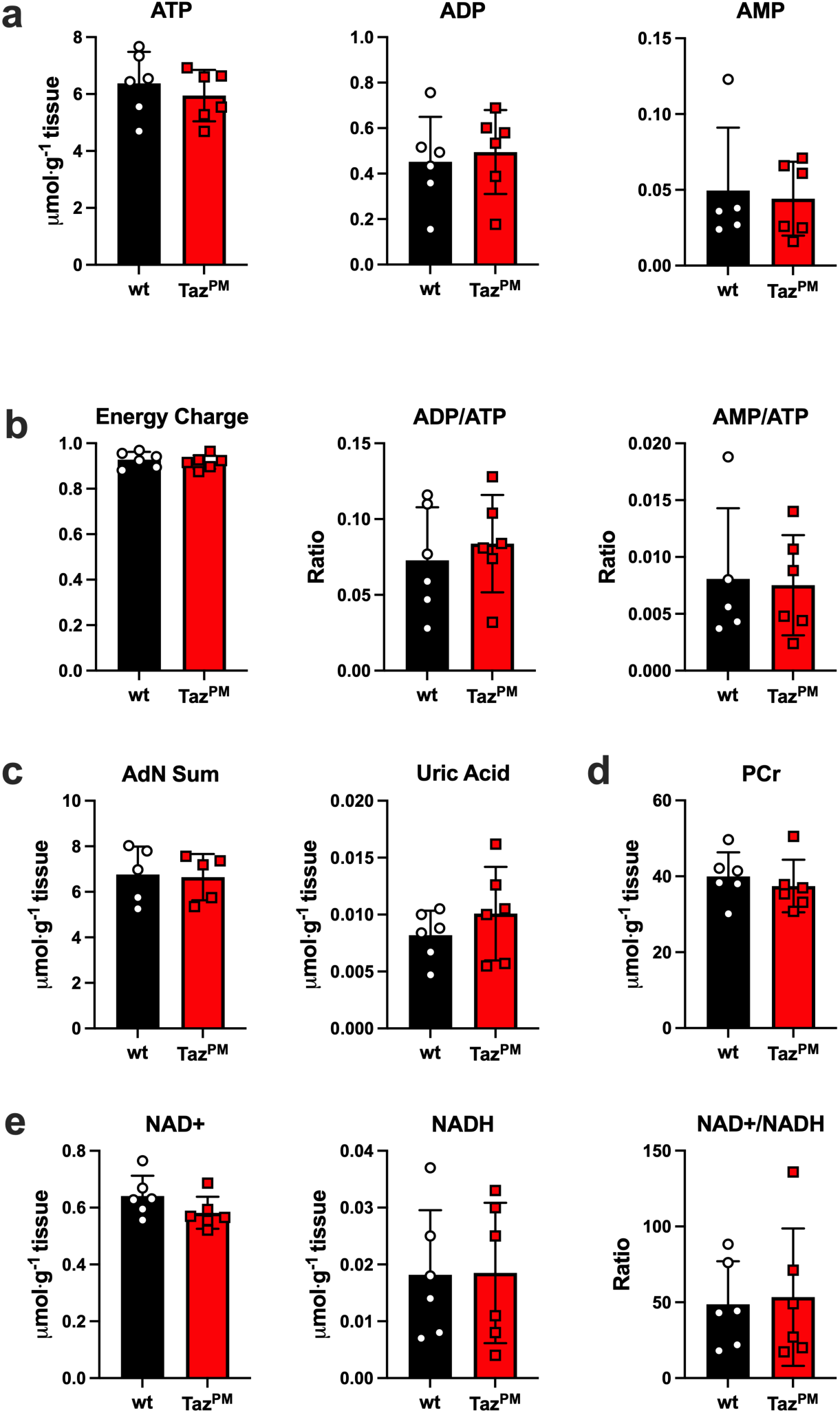
The energetic state in gastrocnemius muscle of *Taz^PM^* mice is the same as that of *wt* mice. (a) Adenine nucleotide (ATP, ADP C AMP) analysis by ultra-performance liquid chromatography (UPLC) of gastrocnemius muscles. (b) Adenine nucleotide calculations of energy charge = (ATP+(0.5*ADP))/(ATP+ADP+AMP), ADP/ATP ratio, and AMP/ATP ratio. (c) Calculation for the sum of adenine nucleotides (AdN Sum = ATP + ADP + AMP) and UPLC analysis of uric acid. (d) UPLC analysis of phosphocreatine (PCr). (e) UPLC analysis of NAD+, NADH, and NAD+/NADH ratio. n=3 per genotype.

Because mitochondrial content and adenine nucleotides differ between muscle fiber types [21, S18], we also measured ATP, ADP, AMP, and PCr in soleus muscles, which are rich in mitochondria and consist predominantly of type 1 and type 2A fibers [S19]. There were no differences in the concentrations of ATP (**Figure S2a**), ADP (**Figure S2b**), AMP (**Figure S2C**), or their sum (**Figure S2d**). Similarly, the calculated energy charge (**Figure 2Se**), the ADP/ATP ratio (**Figure S2f**), the AMP/ATP ratio (**Figure S2g**), and PCr (**Figure S2h**) were also unchanged. Therefore, even in muscles with a greater concentration of mitochondria and a potential greater dependence on mitochondrial respiration, cellular energetics remained unaffected in *Taz^PM^*.

To explore possible signaling pathways that may mediate the effect of muscle weakness in BTHS, we investigated critical stress-responsive proteins. Canonically, various cellular stresses lead to eIF2α-Ser51 phosphorylation that in turn results in a global attenuation of translation but preferential translation of selective transcription factors such as ATF4 or CHOP, termed the Integrated Stress Response (ISR), which includes the Unfolded Protein Response pathway [24]. The ISR can lead to apoptosis, autophagy, or other cellular adaptations that alleviate stress and are thought to be pro-survival in the short-term but pathogenic when responding to severe stress and/or chronic activation.

Therefore, we performed western blots to investigate important indicators of the ISR (**Figure 7a**). Muscles of *Taz^PM^* mice demonstrated a dramatic increase in eIF2a-Ser51 phosphorylation without changes in total eIF2a protein amount (**Figure 7b**), a key indicator of ISR. Further, stress-activated cytoplasmic CHOP and mitochondrial DELE1 protein levels were also robustly increased (**Figure 7b**), indicating ISR activation in response to *Taz^PM^* muscle stress. Additionally, P53, which is also critical for orchestrating cell fate during stress and interacts with the ISR, was highly induced, while MDM2, a primary negative regulator of P53, remained unchanged (**Figure 7c**). Beta-catenin and GSK3, central members of the Wnt signaling pathway that influence the ISR and play a central role in how muscles respond to stress [S20], were both strongly induced in *Taz^PM^* (**Figure 7c**).

**Figure 7.**
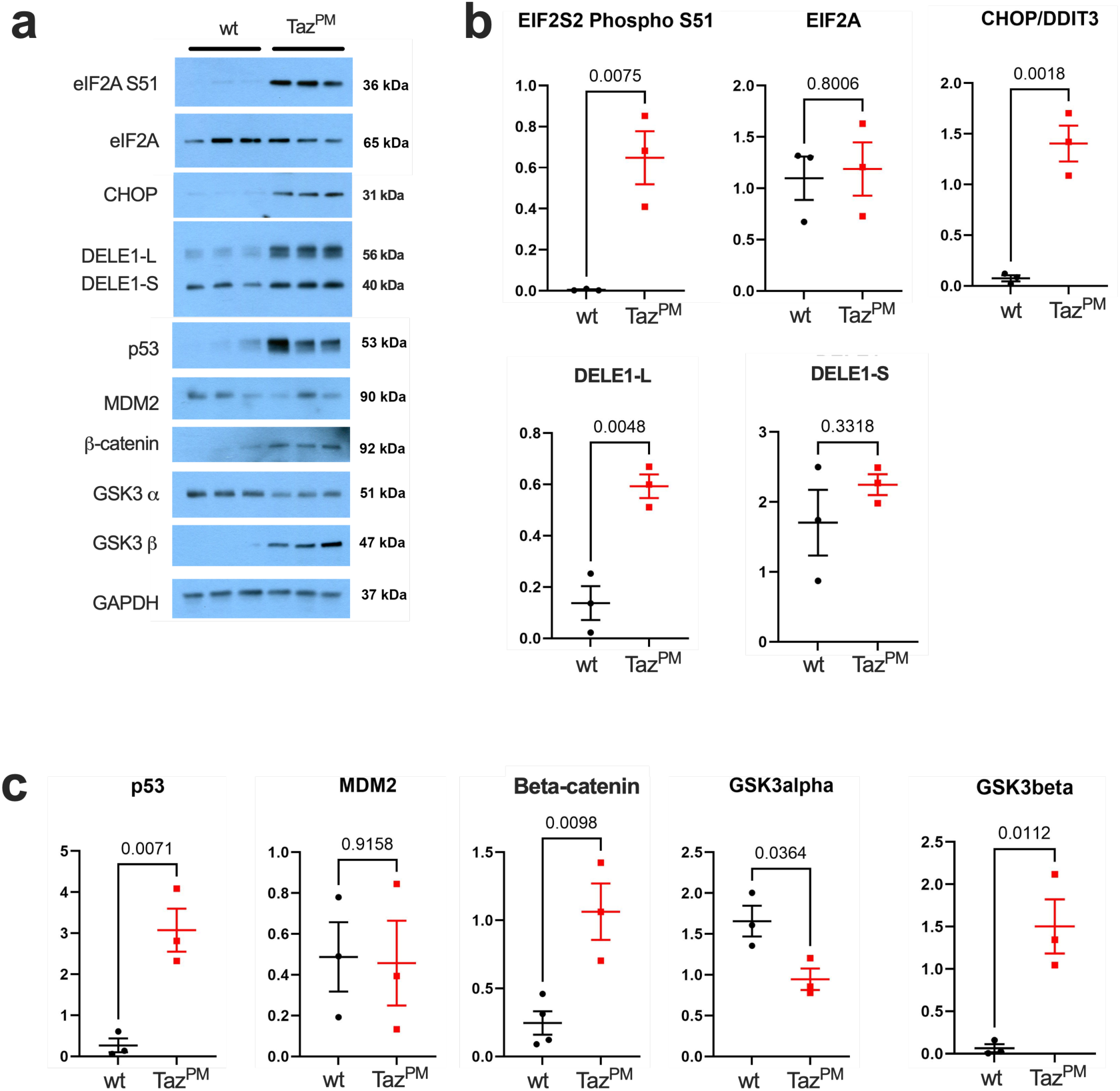
Stress signaling is increased in muscle of *Taz^PM^* mice. (a) Western blots for phosphoSer51-eukaryotic translation initiation factor 2A (eIF2A S51), total eukaryotic translation initiation factor 2A (eIF2A), C/EBP homologous protein (CHOP), Protein Death Ligand Enhancer (DELE1), p53, Mouse double minute 2 homolog (MDM2), β-catenin, Glycogen synthase kinase 3 alpha (GSK3 α), Glycogen synthase kinase 3 beta (GSK3 β), and GAPDH. (b) Quantification of the western blots for ISR proteins eIF2A and CHOP, and mitochondrial stress response DELE1 long form (DELE1-L) and processed form (DELE1-S). (c) Quantification of western blots for p53, MDM2, β-catenin, GSK3 α, and GSK3 β. n=3 per genotype.

Levels of GSK3α protein, which exhibits some overlapping substrate specificity to GSK3β but negatively affects ISR [S21], were significantly reduced in *Taz^PM^* muscle (**Figure 7c**). Furthermore, the number of cells that are Pax7+, a marker of satellite cells necessary for NMJ maintenance [25], is decreased in the *Taz^PM^* muscle (**Figure S3a, b**). As activation of ISR, P53, and Wnt/β-catenin signaling can adversely affect OxPhos, muscle stem cells, muscle regeneration, function, and homeostasis, as well as the NMJ; these findings demonstrate diverse stress-mediated signaling effectors are combinatorially induced to enable adaptation to the chronic loss of mitochondrial transacylase activity in mitochondria of adult *Taz^PM^*.

## Discussion

Muscle weakness and fatigability are cardinal features of BTHS, yet the underlying muscle-specific factors that contribute to this dysfunction remain largely unknown. Using the *Taz^PM^* mouse model with a patient-specific, enzyme-inactivating mutation in Tafazzin, we identified several critical factors that affect muscle contraction performance. First, *Taz^PM^* muscle fibers, regardless of fiber type, are smaller. Smaller muscles, which have less myofibrillar protein, have a decreased capacity to generate force. Second, there is a *Taz^PM^* shift toward a higher percentage of fast type 2B muscle fibers, which have a higher energy cost of contractions and are generally more fatigable [S22]. Third, the number of *Taz^PM^* motor units was decreased, and the expression of NMJ-related proteins was dysregulated, indicating a diminished capacity to fully innervate all muscle fibers. Surprisingly, despite major changes in mitochondrial morphology and mitochondrial protein expression, *Taz^PM^* resting levels of ATP and energy charge were unchanged. However, analysis of signaling proteins suggests that the stress regulated pathways, such as the ISR, are induced in *Taz^PM^* muscles. Therefore, muscle functional changes in BTHS may not be simply due to the direct effects of mitochondrial OxPhos insufficiency or inadequate ATP production but rather an accumulation of neuromuscular remodeling that results in weaker and more fatigable BTHS muscle.

Fiber type predominance is a feature of many mitochondrial myopathies, presenting as either a shift toward slow-twitch type 1 fibers [26] or a shift toward fast-twitch type 2 fibers [27]. Nonetheless, muscle fiber typing has never been reported in BTHS patients or in mouse models of BTHS. In the tibialis anterior muscles of our *Taz^PM^* mouse, we demonstrate a significant shift away from type 2A and 2X fibers toward the type 2B fiber type. The type 2B MHC isoform has the fastest contraction speed but also has the highest energetic demand per contraction of typical adult muscle myosin [S23, S24]. Therefore, this shift to a more energetically demanding fiber type would make any amount of contraction more demanding and likely contribute to muscular fatigue.

Taffazin deficiency leads to consistent defects in mitochondrial respiration in skeletal muscle [6, 10, S2]. To investigate whether this affects resting cellular energetics, we measured adenine nucleotide (ATP, ADP, and AMP) and phosphocreatine levels.

Perhaps surprisingly, these levels were unchanged in the current study in both the predominantly glycolytic, fast-twitch gastrocnemius muscle and the predominantly oxidative, mixed-fiber (∼30% type 1, ∼70% type 2 fibers [S19]) soleus muscle, demonstrating that ATP production is entirely sufficient to meet basal energetic demands in muscles that have different mitochondrial contents. This apparent disassociation between impaired mitochondrial function and normal adenine nucleotide levels has been noted in other mitochondrial myopathies [17, 21]. Regardless, we do not interpret these findings as indicating that energetics have no role in the BTHS phenotype. Our measures were performed on non-contracting muscle tissue, where energy demand is relatively low. During muscle contraction, ATP demand can increase more than >100-fold, far exceeding resting needs. Therefore, ATP production limitations during or immediately after contractions may be an important feature of BTHS, as previously suggested [10]. Future studies are required to determine the extent to which mitochondrial and/or glycolytic ATP production becomes limiting at different contraction intensities and durations.

Muscle force production and fatigability are dependent on effective innervation of muscle fibers. In the hindlimb muscles of *Taz^PM^*, we observed a 60% decrease in the MUNE, accompanied by an increase in SMUP. This loss of motor units is entirely consistent with our observed muscle fiber type grouping, which is an indicator of motor neuron denervation and reinnervation [S25]. Importantly, loss of motor unit number is found with other conditions of muscle weakness, such as ALS, spinal muscle atrophy, generalized neuropathies, and aging [28]. Despite the substantial changes in MUNE in the *Taz^PM^*, CMAP was relatively unchanged. Because CMAP is the summed output from all motor units, a normal CMAP suggests that all (or nearly all) muscle fibers are innervated. However, experimentally, CMAP is elicited by supramaximal stimulation of the peripheral nerve, leaving open the possibility that physiological innervation of some fibers may still be impaired. Regardless, our electrophysiological data clearly demonstrate extensive motor unit remodeling in *Taz^PM^*, with fewer surviving motor neurons innervating a greater number of muscle fibers. Since the larger, fast-type motor units are more fatigable [29], this remodeling likely contributes to the greater fatigability observed in BTHS. Further, if motor unit remodeling manifests early in the disease, a dramatically lower number of motor units may contribute to delayed motor development and difficulties with fine motor skills.

Many mitochondrial disorders are associated with NMJ defects [30]. The remodeling of motor units may originate from either the presynaptic nerve terminal or postsynaptic motor endplate side of the NMJ. Supporting a presynaptic origin, both Pax7- positive cells (satellite cells) and muscle-specific mitochondrial function are critical for NMJ stability [S26]. On the other hand, mitochondria are enriched on the presynaptic side of the NMJ, where they serve in specialized roles in ATP provision, Ca^2+^ buffering, and even axonal protein synthesis [31]. Unlike most CNS synapses, which tend to rely heavily on glycolysis, NMJs are almost entirely dependent on OxPhos for ATP supply [31]. This makes the presynaptic NMJ especially vulnerable to alterations in cristae organization, OxPhos supercomplex stability, and Ca^2+^ buffering, all of which are impaired via TAFAZZIN deficiency [S27]. Thus, TAFAZZIN-related mitochondrial dysfunction could compromise NMJ stability from either side of the NMJ.

Consistent with overall remodeling of motor units, we observed altered expression of proteins associated with the postsynaptic region. Specifically, DOK7 and MUSK were decreased, and CHRNA1 was increased. For proteins associated with the presynaptic region, levels of SNAP-25, α tubulin, and β tubulin were unchanged. Taken together, these findings suggest a remodeling of the NMJ and a potential disruption of communication between motor neurons and muscle fibers. Specific NMJ functionality testing, by comparing muscle force production via nerve stimulation versus direct muscle stimulation (e.g., [S28]), will be required to test this hypothesis directly.

The ISR is a conserved signaling network that allows cellular adaptation to various stressors by inhibiting global translation while promoting translation of select proteins [S29]. *Taz^PM^* skeletal muscle demonstrates canonical activation of the ISR characterized by large increases in eIF2A Ser51 phosphorylation, accompanied by increased expression of central ISR-associated CHOP and DELE1 proteins. This mirrors the induction of ISR markers that have been noted in hearts of other BTHS models [16, S30]. The downstream stress-responsive transcription factor CHOP plays a major role in tuning ISR duration and cellular adaptation to mitochondrial stress [32], whereas upstream DELE1 is a mitochondrial protein that senses disruption of the inner mitochondrial membrane [33] that can drive eIF2A Ser51 phosphorylation and activate the ISR. The finding that full-length mitochondrial DELE1 robustly accumulates but that cleaved cytoplasmic DELE1 is not significantly altered suggests an intact cytoplasmic ISR that may be adaptively beneficial in counteracting *Taz^PM^* inner mitochondrial membrane dysfunction [34]. Further, p53 is overexpressed in *Taz^PM^* and likely modifies the ISR via protecting adenine nucleotide levels and may underlie the reduced fiber size and central nuclei [35, 36]. Additionally, stress-related proteins GSK3β and β-catenin proteins play opposite roles in skeletal muscle health, and mitochondrial functions were increased in *Taz^PM^*. While GSK3β and β-catenin are not canonical ISR proteins, their altered balance likely helps increase muscle fiber protein breakdown, promotes a more glycolytic phenotype, inhibits mitochondrial biogenesis pathways, and plays a significant role in controlling mitochondrial content and function [S20, 37, 38].

Previous knockdown (KD) and knockout (KO) mouse models have been invaluable for establishing TAFAZZIN’s role in CL remodeling. However, they are severely limited for dissecting the pathogenic mechanisms of skeletal muscle and NMJ. KD and KO models exhibit substantial embryonic/perinatal lethality, constraining adult phenotyping [39]. This is true even in tissue-restricted KO models, which are dominated by severe cardiac dysfunction. In contrast, the D75H point mutant reproduces a clinically observed BTHS mutation that maintains normal/near-normal TAFAZZIN protein abundance but severely impairs enzymatic activity, reflecting the genotype-phenotype configuration seen in a BTHS patient [18]. Although the D75H (239-2A>G) mutation seems to be exclusive to this patient, additional pathogenic substitutions at the same residue (D75N, D75R) and in adjacent positions (D74E, P76R, P76L), variants of unknown significance (238+6C>T, 238+5C>T, 239-3T>A) and benign variants (238+10C>A, 238+11C>G, 239-6delC, 239-6C>T, 293-3T>A) have also been independently reported in male BTHS patients [S1]. Asp75 lies immediately adjacent to the conserved HX_4_D active motif, which is essential for acyltransferase activity. This proximity situates Asp75 within a structurally and functionally sensitive region, where clustering of pathogenic variants there underscores its critical importance for TAFAZZIN enzymatic function. Many BTHS patients present with missense variants rather than complete deletions, and therefore exhibit residual TAFAZZIN protein. The configuration of our model enables the separation of protein quantity from enzyme function, allowing us to uncover muscle-specific and NMJ-level pathophysiology (fiber size/type remodeling, clumping, reduced MUNE, altered NMJ proteins) that would have likely been masked in null backgrounds. Finally, patient-derived models are exceptionally well-suited for testing therapies that stabilize or enhance mutant TAFAZZIN activity, such as gene therapy, pharmacological chaperones, or CL-targeted agents (e.g., elamipretide [40]).

In summary, deficiency of Tafazzin enzyme activity results in widespread neuromuscular remodeling in adult mice. Muscle fibers are smaller, fiber type shifts to the fastest type 2B, motor unit number is decreased, neuromuscular junction protein amounts are dysregulated, and robust stress signaling is active. Together, these factors explain, at least in part, the reduced force production and increased fatigability in BTHS. However, cellular energetics, as defined by resting levels and ratios of adenine nucleotides and PCr, remain unchanged by the loss of Tafazzin enzymatic activity and associated reduced CL levels. Therefore, muscle functional changes in BTHS are not solely due to the direct effects of mitochondrial insufficiency but rather an accumulation of widespread neuromuscular remodeling and stress signaling pathways activation.

## Grant Funding

This work was supported in part with financial support from the NIH R01 HL159436 (to SJC) and R37CA299888 (to JRH), the Barth Syndrome Foundation (to SJC), the Hevolution Foundation HF-GRO-23-1199172-46 (to JJB), the American Cancer Society RSG-24-1258280 (to JRH), and the Herman B Wells Center in part from the Riley’s Children’s Foundation (to SJC & JJB).

## Acknowledgements

The authors would like to thank Dr. Brian Piechala and Ms. Emily Yang for assistance with neuromuscular junction staining. We also thank the Indiana University School of Medicine Center for Electron Microscopy for technical assistance.

## Notes

### Competing Interest Statement

The authors have declared no competing interest.

