## Supplementary Information for "Deficient Cardiolipin Remodeling Alters Muscle Fiber Composition and Neuromuscular Connectivity in Barth Syndrome"

Methods

- Muscle processing for immunofluorescence staining
- Fiber Typing and centrally nucleated fibers
- Neuromuscular Junction Imaging
- Ultra-performance liquid chromatography

Tables

- Table S1: Primary antibodies
- Table S2: Secondary antibodies

Figures

- Figure S1. Central nuclei are decreased in gastrocnemius muscles of $Taz^{PM}$ mice.
- Figure S2. Adenine nucleotide concentrations are unchanged in soleus muscles of $Taz^{PM}$.
- Figure S3. The number of Pax7 pos cells per fiber is decreased in $Taz^{PM}$ muscle.

References

**Methods**

*Muscle processing for immunofluorescence staining*

Tibialis anterior (TA) and gastrocnemius muscles were excised, frozen in isopentane cooled in liquid nitrogen, and then stored at -80°C until further processing. Muscles were sectioned at -21°C using a Leica CM1950 cryostat to obtain 10 μm transverse sections of the TA for fiber typing analysis and SDH histochemistry, or 40 μm longitudinal sections of the gastrocnemius for neuromuscular junction (NMJ) imaging. Sections were mounted on microscope slides (48311-703, VWR). Slides were stored at -20°C.

*Fiber typing and centrally nucleated fibers*

Slides with transverse sections were permeabilized with 1X phosphate-buffered saline (PBS) + 0.1% Triton X-100 and blocked with a solution of 1X PBS + 0.5% BSA (BAC62, Equitech-Bio) + 10% goat serum (PCN5000, Thermo Fisher Scientific). Primary antibodies for Type I myosin heavy chain (MHC), Type IIA MHC, Type IIB MHC and laminin (Table S1) were diluted together in a 0.5% BSA + 2% goat serum + 1X PBS solution and incubated overnight in a humidified chamber at 4°C. After washing with 1X PBS, the slides were incubated for one hour with the following secondary antibodies (Table S2). Post-incubation, slides were washed 3 times with 1X PBS for 5 minutes each.

For centrally nucleated fiber analysis, a slide probed only with laminin was incubated at room temperature in the dark in a 4',6-Diamidino-2-phenylindole dihydrochloride, 2-(4-Amidinophenyl)-6-indolecarbamidine dihydrochloride (DAPI) (D9542,

MilliporeSigma) solution for 15 minutes, repeating the PBS washes after incubation. Slides were mounted using SlowFade Diamond Antifade Mountant (S36963, Thermo Fisher Scientific) and sealed.

Images were acquired with a Keyence BZ-X800 fluorescence microscope (Keyence Corp) at the Indiana Center for Biological Microscopy using a 10x objective. Initial image stitching and deconvolution were done in the BZ-X800 Analyzer software. The resulting images were analyzed using QuantiMus, a Flika plugin that allows for measuring the cross-sectional area (CSA), quantifying centrally nucleated fibers (CNF), and assessing the fluorescence intensity of individual myofibers.

*Neuromuscular junction imaging*

Slides with longitudinal 40-μm triceps surae tissue sections were fixed in 1% PFA (15710, Electron Microscopy Sciences) for 30 minutes and then washed with 1X PBS three times. Slides were blocked with a solution of 1X PBS + 0.3% Triton X-100 (X100, Millipore Sigma) + 1% BSA (BAC62, Equitech-Bio) + 10% donkey serum (50-588-37, Millipore Sigma) + M.O.M (MKB-2213-1, Vector Laboratories) for one hour to prevent nonspecific antibody binding. Primary antibodies (Table S1) were diluted in a 1% BSA + 0.3% Triton-X + 1X PBS solution and then incubated overnight in a humidified chamber at 4°C. After incubation, the slides were washed in 1X PBS and incubated for two hours with α-Bungarotoxin, CF@488A (00005, Biotium) at 1:200, along with the following secondary antibodies (Table S2). Post-incubation, slides were washed with 1X PBS, mounted using SlowFade Diamond Antifade

Mountant (S36963, Thermo Fisher Scientific), and sealed. All images were acquired within three days of probing.

NMJ images were acquired using a Leica SP8 Lightning confocal microscope (Leica Microsystems) equipped with a 63x glycerol immersion objective. Z-stacks were collected at 2 μm intervals, covering a total depth of 20–30 μm. Initial image stitching, deconvolution, and maximum projections were done in LAS X software. Any subsequent image processing was carried out using ImageJ.

*Ultra Performance Liquid Chromatography*

To maintain nutritive blood flow to hindlimb muscles, mice were deeply anesthetized with 2% isoflurane in oxygen delivered via a nosecone and placed on a 37 °C heating pad. Gastrocnemius and soleus muscles were excised and immediately freeze-clamped using liquid nitrogen-cooled steel clamps. Rapid freezing is essential for metabolic quenching and preservation of labile high energy phosphates. Frozen muscles were weighed to the nearest 0.1 mg. Muscles were homogenized with glass tubes and glass pestles (Kontes) at a ratio of 1 mg muscle per 49 μL extraction solution (80 % UPLC-grade methanol: 20% ultrapure water). Homogenized samples were incubated at -20°C for 30 minutes, then centrifuged at 15,000 x g for 10 min at 10°C. The pellets were discarded, and the supernatant was transferred to pre-chilled tubes.

Analytes were separated and measured using a Waters Acquity Premier UPLC system, Acquity Premier Tunable UV Detector, QDa Mass Detector, and Acquity Premier HSS T3 column with 1.8 μm VanGuard fit 2.1 × 150 mm (p/n 186009472, Waters).

Chromatography buffers and conditions were developed previously to elicit baseline resolution [33, 41], which allows quantification by UV absorbance and avoids matrix effects errors. Phosphocreatine was quantified by absorbance at 210nm. Uric acid was quantified at 290nm. NADH was quantified at 338nm. NAD+, ATP, ADP, AMP, and IMP were quantified at 254nm. Identity was confirmed by mass detection with Waters QDa.

**Table S1. Primary antibodies**

| Antibody Name | Host species | Source | Catalog number | Dilution | Application |
|---|---|---|---|---|---|
| Laminin | Rabbit | Sigma-Aldrich | L9393 | 1:300 | IF |
| Myosin Heavy Chain 1 | Mouse | DSHB | BA-F8 | 1:100 | IF |
| Myosin Heavy Chain 2A | Mouse | DSHB | SC-71 | 1:100 | IF |
| Myosin Heavy Chain 2B | Mouse | DSHB | BF-F3 | 1:100 | IF |
| Synapsin-1 | Rabbit | Cell Signaling | 5297 | 1:200 | IF |
| $\beta$-Tubulin III | Mouse | Sigma-Aldrich | T8578 | 1:200 | IF |
| Pax7 | Mouse | DSHB | PAX7 | 1:100 | IF |
| $\alpha$ tubulin | Mouse | Sigma-Aldrich | T5168 | 1:1000 | WB |
| ATP5a1 | Rabbit | Proteintech | 14676-1-AP | 1:10,000 | WB |
| $\beta$-catenin | Rabbit | Abcam | ab32572 | 1:30,000 | WB |
| $\beta$ tubulin | Rabbit | Abcam | ab18207 | 1:5,000 | WB |
| CHRNA1 | Rabbit | Abcam | ab308306 | 1:5,000 | WB |
| Citrate synthase | Rabbit | Cell Signaling | 14309 | 1:5,000 | WB |
| DELE1 | Mouse | Santa Cruz | Sc-515080 | 1:1,600 | WB |
| DOK7 | Rabbit | Abcam | ab75049 | 1:400 | WB |
| eIF2A, phospho-S51 | Rabbit | Abcam | ab32157 | 1:2,000 | WB |
| eIF2A, total | Rabbit | Abcam | ab169528 | 1:2,500 | WB |
| GAPDH | Mouse | Sigma-Aldrich | G8795 | 1:25,000 | WB |
| GSK3 $\alpha$ | Rabbit | Abcam | ab40870 | 1:30,000 | WB |
| GSK3 $\beta$ | Rabbit | Abcam | ab32391 | 1:1,000 | WB |
| MCU | Rabbit | Cell Signaling | 14997 | 1:650 | WB |
| MDM2 | Rabbit | Bio-Rad | AHP1329 | 1:3,000 | WB |
| MUSK | Rabbit | Invitrogen | PA1-1741 | 1:400 | WB |
| NDUFB8 | Mouse | Abcam | ab110242 | 1:1,000 | WB |
| p53, total | Mouse | Santa Cruz | sc-71820 | 1:1,250 | WB |
| Rapsyn | Rabbit | Abcam | ab156002 | 1:5,000 | WB |
| SDHA | Mouse | Abcam | ab14715 | 1:30,000 | WB |
| SNAP25 | Rabbit | Abcam | ab109105 | 1:1,000 | WB |
| TMEM65 | Rabbit | Abcam | ab236861 | 1:2,500 | WB |
| VDAC1 | Rabbit | Invitrogen | PA1-954A | 1:80,000 | WB |

**Table S2. Secondary antibodies**

| Antibody Name | Host species | Source | Catalog number | Dilution | Application |
|---|---|---|---|---|---|
| Anti-Rabbit IgG (H+L) | Goat | Invitrogen | A-21244 | 1:500 | IF |
| Anti-Mouse IgG2b | Goat | Invitrogen | A-21140 | 1:500 | IF |
| Anti-Mouse IgG1 | Goat | Invitrogen | A-21121 | 1:500 | IF |
| Anti-Mouse IgM | Goat | Invitrogen | A-21045 | 1:500 | IF |
| Anti-Mouse IgG2a | Goat | Invitrogen | A-21143 | 1:500 | IF |
| Anti-Rabbit IgG (H+L) | Goat | Bio-Rad | 170-6515 | 1:8,000 | WB |
| Anti-Mouse IgG (H+L) | Goat | Jackson ImmunoResearch | 115-035-146 | 1:8,000 | WB |

DSHB=Developmental Studies Hybridoma Bank, IF=immunofluorescence, WB=western blot

### Figure S1

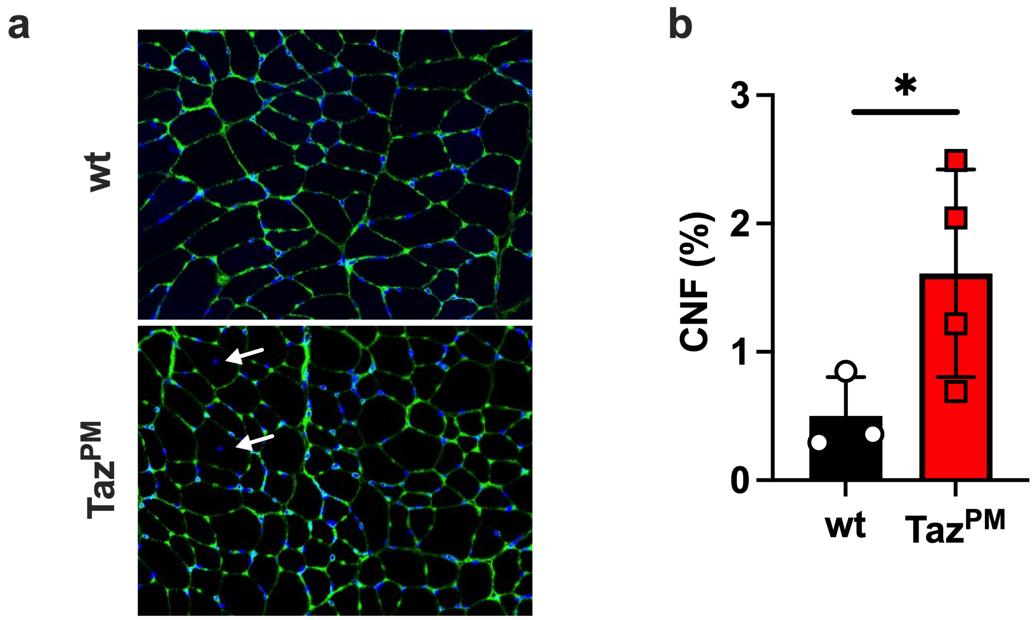


**Figure S1.** (a) Representative immunofluorescent images of gastrocnemius muscle stained for laminin (green) and nuclei (blue) in *wt* (Top) and *Taz*$^{PM}$ mice. White arrows point to central nuclei. (b) Quantification of centrally nucleated fibers is calculated as percent of fibers. n=3-4 per genotype. *p<0.05

### Figure S2

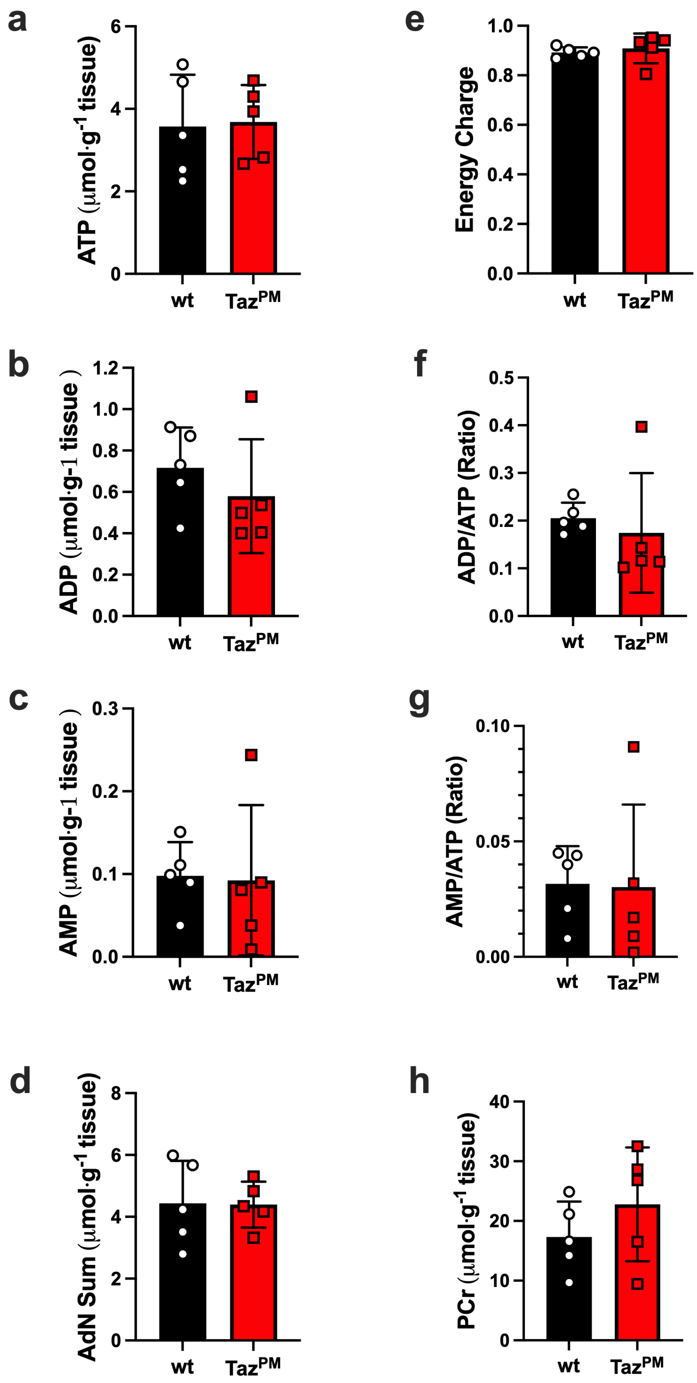

a
ATP (μmol·g-1 tissue)
6
4
2
0
wt
TazPM
e
Energy Charge
1.0
0.8
0.6
0.4
0.2
0.0
wt
TazPM
b
ADP (μmol·g-1 tissue )
1.2
1.0
0.8
0.6
0.4
0.2
0.0
wt
TazPM
f
ADP/ATP (Ratio)
0.5
0.4
0.3
0.2
0.1
0.0
wt
TazPM
c
AMP (μmol·g-1 tissue )
0.3
0.2
0.1
0.0
wt
TazPM
g
AMP/ATP (Ratio)
0.10
0.05
0.00
wt
TazPM
d
AdN Sum (μmol·g-1 tissue)
8
6
4
2
0
wt
TazPM
h
PCr (μmol·g-1 tissue)
40
30
20
10
0
wt
TazPM

**Figure S2. Adenine nucleotide concentrations are unchanged in soleus muscles of *Taz$^{PM}$*.** Ultra-performance liquid chromatography analysis of (a) ATP, (b) ADP, and (c) AMP in soleus muscles of *wt* and *Taz$^{PM}$* mice. Calculations of (d) AdN Sum = ATP + ADP + AMP, (e) Energy Charge = (ATP + (0.5 * ADP)) / (ATP + ADP + AMP), (f) ADP/ATP ratio, (g) AMP/ATP ratio, and (h) phosphocreatine (PCr). n=5 per genotype.

### Figure S3

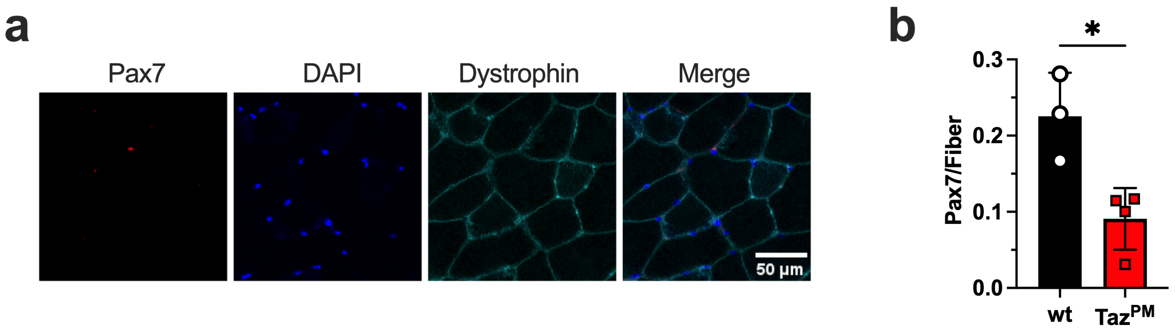


**Figure S3. The number of Pax7 positive cells per fiber is decreased in *Taz$^{PM}$* muscle.** (a) Representative immunofluorescent images of gastrocnemius muscle stained for dystrophin (green), nuclei (blue), and Pax7 in *wt* mice. (b) Quantification of Pax7 positive cells per number of fibers. n=3-4 per genotype. *$p<0.05$
